# PARP mediated chromatin unfolding is coupled to long-range enhancer activation

**DOI:** 10.1101/155325

**Authors:** Nezha S. Benabdallah, Iain Williamson, Robert S. Illingworth, Shelagh Boyle, Graeme R. Grimes, Pierre Therizols, Wendy A. Bickmore

## Abstract

Enhancers are critical regulators of gene expression and can be located far from their target gene. It is widely assumed that mechanisms of enhancer action involve reorganization of three-dimensional chromatin architecture, but this is poorly understood. Here we identify a novel mechanism of long-range enhancer associated chromatin reorganization. At the Sonic hedgehog (*Shh*) locus we observe large-scale decompaction of chromatin between *Shh* and its brain enhancers in neural progenitor cells. We show that the chromatin unfolding is dependent on activation of the enhancers, not the promoter, is impeded by chromatin-bound proteins located between the enhancer and promoter, and is mediated by the recruitment of Poly (ADP-Ribose) Polymerase 1. We suggest that large-scale chromatin decompaction, analogous to the inducible puffs in *Drosophila* polytene chromosomes, represents a new mechanism of chromatin reorganization coupled to long-range gene activation from mammalian enhancers and that seems incompatible with a chromatin-looping model of enhancer-promoter communication

## Introduction

Enhancers are cis-regulatory sequences, often located within the non-coding portion of the genome, which function to tightly regulate spatial and temporal gene expression in development and physiology. Enhancers can operate when located proximal to, or very distant (100s to 1000s of kb) from, their target gene (Vernimmen and Bickmore, 2015). Well established molecular signatures of functional enhancers include; clustered sequence-specific transcription factor binding sites and DNaseI hypersensitive sites (DHS), specific histone modifications e.g. H3K4me1 and acetylation of specific lysine residues on histone H3 (H3K27ac, H3K64ac, H3K122ac) and H4 (H4K16ac) and, in some cases, eRNA transcription (Kim et al., 2010; Pradeepa et al., 2016; Shlyueva et al., 2014; Taylor et al., 2013). However, less is known about the mechanisms by which enhancers communicate with, and control the expression of, their target gene promoter(s).

For proximal enhancers, it has been proposed that activation signals nucleated by bound transcription factors (TFs) then spread or move towards the target gene, by reorganising or modifying the intervening chromatin (Benabdallah and Bickmore, 2015; Engel et al., 2008; Hatzis and Talianidis, 2002; Wang et al., 2005; Zhao and Dean, 2004; Zhu et al., 2007). These ‘tracking’ or ‘facilitated-tracking’ models are compatible with enhancers located in the vicinity of their target promoter, but have been considered unlikely as a mechanism for more distal enhancers. For very long-range regulation, direct communication between the enhancer and the promoter is thought to occur through interaction of protein complexes bound at both sites, with looping-out of the intervening chromatin. Chromatin ‘looping’ has best been illustrated for interactions between the β-globin gene and its locus control region (LCR): enhancer-promoter interactions have been detected by chromosome conformation capture (3C) methods (Carter et al., 2002; Tolhuis et al., 2002) and experimentally forced enhancer-promoter chromatin looping contributes to transcriptional activation (Bartman et al., 2016; Deng et al., 2012, Deng et al., 2014). Fluorescence in situ hybridisation (FISH) has also been used to visualise the spatial juxtaposition of a target gene (*Shh*) with its distant (1Mb) limb enhancer, with a looping-out of the intervening chromatin, specifically in *Shh* expressing tissues of the developing limb bud (Williamson et al., 2016). A general compaction of the chromatin fibre between enhancer and promoter, rather than a discrete loop, has also been suggested (Williamson et al., 2012, 2014) and this may facilitate factors recruited at enhancers finding their promoter-proximal binding sites through diffusional mechanisms (Benabdallah and Bickmore, 2015). However, to date, no other specific chromatin conformation has been demonstrated to contribute to long-range gene regulation from enhancers.

The sonic hedgehog morphogen (Shh) governs the growth and patterning of many tissues during development. The precise spatial and temporal control of *Shh* expression is regulated by tissue-specific enhancers located; within the introns of the gene, upstream of the *Shh* transcription start site (TSS) in a large (750kb) gene desert, and within genes at the far end of the gene desert (Anderson and Hill. 2014). *Shh* expression is important for several aspects of brain development. Shh-Brain-Enhancers-5 (SBE5), SBE2/3, SBE4 and SBE6 have been demonstrated to drive expression in the midbrain and anterior domains of the developing brain and are located 780, 450, 350 and 100 kb upstream of the *Shh* TSS, respectively (Jeong, 2006; Benabdallah et al., 2016; Yao et al., 2016). As distal enhancers, the SBE elements might be expected to loop into spatial proximity of *Shh* during its activation in neural progenitors cells (NPCs). However, here we show that induction of *Shh* expression in NPCs, or by synthetic enhancer activation in mouse embryonic stem cells (ESCs) does not lead to detectable co-localisation between *Shh* and SBE6, SBE4, or SBE2/3. Rather, we show that enhancer activation leads to an unfolding of chromatin between *Shh* and SBE4/SBE6 that appears to be mediated by the recruitment and activity of Poly (ADP-Ribose) Polymerase 1.

## Results

### Increased nuclear separation of Shh and Shh-Brain-Enhancers upon neural differentiation

To analyse the spatial relationship of *Shh* with its known brain enhancers in the nucleus we differentiated 46c ESCs (Ying et al., 2003) into NPCs. Efficient differentiation was monitored through *Sox1*-GFP fluorescence (Benabdallah et al., 2016).

As the known *Shh*-Brain-Enhancers SBE5, SBE2/3, SBE4 and SBE6 are located 780, 450, 350 and 100 kb upstream of *Shh*, respectively (Figure 1A), the favoured mechanism by which these enhancers control *Shh* would involve a physical looping between them and the *Shh* promoter. To test this we used super-resolution microscopy, in conjunction with 3D fluorescence in situ hybridisation (3D-FISH), to visualise the spatial proximity of the enhancers and *Shh* before and after neural differentiation (Williamson et al., 2016). The squared distances between two FISH probes typically have a linear relationship to the genomic distance that separates them (Gilbert et al., 2004; van den Engh et al., 1992), but is also influenced by chromatin folding (Eskeland et al., 2010; Williamson et al., 2014). Consistent with enhancer-promoter chromosome looping, we have previously demonstrated that spatial juxtaposition of *Shh* and it ZRS limb enhancer, with displacement of an intervening genomic region, is restricted to the time and place of *Shh* expression in the developing limb (Williamson et al., 2016). If chromosome looping also occurs between the brain enhancers and *Shh* upon neuronal activation we anticipated that hybridisation signals for the SBE distal enhancers would display a higher frequency of co-localisation with *Shh (*inter-probe distances ≤ 0.2 μm), and shorter average inter-probe distances, in NPCs as compared with undifferentiated ESCs.

**Figure 1.**
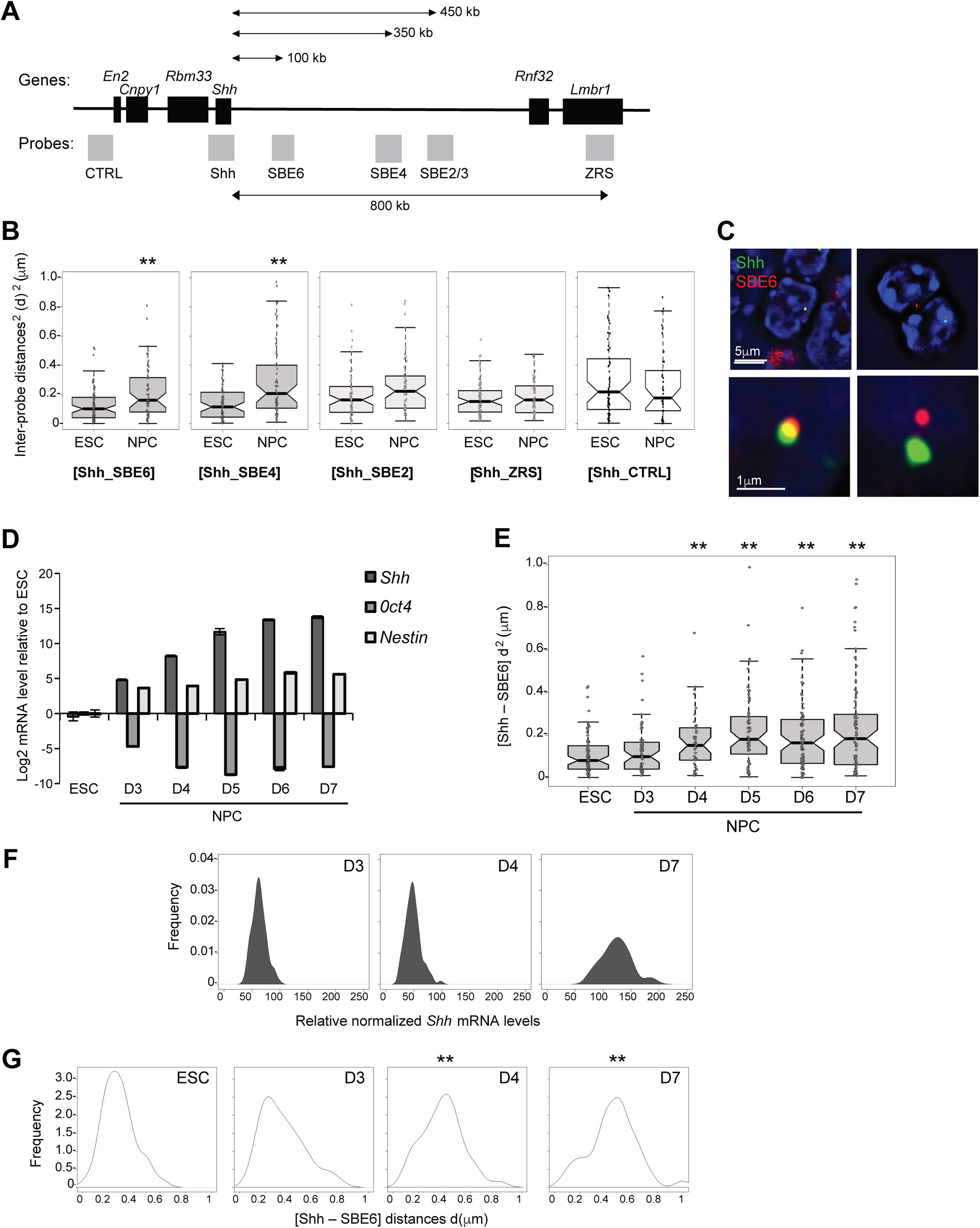
Shh-Brain-Enhancer chromatin decompaction during neuronal differentiation. A) Map of *Shh* regulatory domain showing the genes (black boxes) and fosmid FISH probes (grey boxes). B) Boxplots showing the distribution of squared interprobe distances (d^2^) in pm between *Shh* and SBE6, SBE4, SBE2/3, ZRS and CTRL probes in the nuclei of ESCs and NPCs. Box indicates interquartile range of the median (25 to 75 percentile) with the notch displaying a 95% confidence interval around the median. ** p <0.01. C) 3D-SIM image illustrating *Shh*-SBE6 separation in NPCs after seven days of differentiation from ESCs. Scales bars are 5μm (top) and 1μm (bottom). D) *Shh*, Oct4 and *Nestin* expression assayed by qRT-PCR during a time course of NPC differentiation. Graph shows mean (± SEM) log2 mRNA levels relative to *Gapdh* and normalized to the level in ESC for one biological replicate (three technical replicates). E) Boxplots showing *Shh*-SBE6 squared interprobe distances in cell populations corresponding to the expression data in D. F) Kernel density plots showing *Shh* mRNA expression of single NPCs relative to *Gapdh* and normalised to expression in ESC. G) Kernel density plots showing the distribution of Shh-SBE6 FISH distances (μm) in undifferentiated ESCs and at days 3, 4 and 7 (D3, D4, D7) of differentiation. Density is an arbitrary unit based on the frequency of the occurrence, and the total counts and the size of the population i.e. the binning of the data. Statistical data relating to FISH data from this figure and replicate experiments are in Table S1.

We performed 3D FISH on 46c ESCs that do not express *Shh*, and on NPCs obtained after seven days of differentiation, when *Shh* is expressed (Benabdallah et al., 2016) and we imaged the slides by 3D-Structured Illumination Microscopy (3D-SIM). Surprisingly, upon *Shh* activation there was a significant increase in inter-probe distances between *Shh* and both the SBE6 and SBE4, located 100 and 350 kb 5’ of *Shh* respectively (Figures 1B and 1C). Inter-probe distances between *Shh* and the more distant SBE2/3 or ZRS were not significantly changed, nor were those to a control probe (CTRL) located 340kb downstream of *Shh* and not within the *Shh* regulatory region (Figures 1A and 1B, Table S1). This suggests that a large-scale chromatin unfolding occurs upon *Shh* neural induction and that this is limited to the region 300kb 5’ of the *Shh* TSS. It is noteworthy that, within this 300kb region, the most prominent peaks of H3K3me1 and H3K27ac gained during this NPC differentiation programme occur at SBE6, and that SBE6 is required for full induction of *Shh* during this differentiation programme (Benabdallah et al., 2016).

The increased separation between *Shh* and SBE6/SBE4 upon *Shh* induction does not seem compatible with the formation of a chromatin loop between *Shh* and these two neural enhancers. To assess if ‘looping’ occurs at an earlier time point, we analyzed *Shh* expression at earlier stages during the NPC differentiation time course (days 3 to 7). *Shh* expression initiated on day 3 (D3) and increased steadily until D6 or D7 (Figure 1D). In all replicate experiments, *Shh*-SBE6 nuclear distances were significantly increased from D4 onwards (Figure 1E). Though *Shh*-SBE6 distances were somewhat increased at D3 this did not reach statistical significance in this experiment, but in other replicates, there was a significant Shh-SBE6 distance increase by D3 (Figure S1A). These data support the notion that no stable chromatin loop structures exist between SBE6 and *Shh* throughout this neural differentiation programme (Table S1).

To determine whether *Shh* activation is coupled with the observed increased promoter-enhancer separation across the cell population, or whether there is a sub-population of NPCs that express *Shh* at high levels with a looped chromatin conformation (short inter-probe distances) that are obscured in the whole population, we performed single cell qRT-PCR on cells at D3, D4 and D7 of NPC differentiation. These data show that in most cells of the population there is increased *Shh* expression at D3 and D4, (Figures 1F & S1B), and even higher levels of *Shh* expression by D7, consistent with the cell population averaged expression data (Figure 1D). For the same cell populations, *Shh* - SBE6 inter-probe distances start to increase at D3 and shift homogenously towards greater distances at D4 and D7 (Figures 1G), mirroring the population wide increase in *Shh* expression. Across this time course, there was a corresponding decrease in the frequency of alleles with Shh-SBE6 inter-probe distances (≤200nm) that we would consider compatible with enhancer-promoter juxtaposition (Figures 1G and S1C) (Williamson et al., 2016).

To show that 3D FISH analysis is capable of detecting a chromosome loop in ESCs, we created an artificial Shh-SBE interaction. Targeted tethering (using zinc fingers) of the self-association domain (SA) of LIM domain-binding protein 1 (LDB1) has been used previously to force a chromatin loop at the β-globin locus (Bartman et al., 2016; Deng et al., 2012, 2014). Using a similar approach, but employing transcription activator-like effector (Tale) proteins to direct site-specific binding (Therizols et al., 2014), we tethered the LBD1 SA to the *Shh* promoter (tShh-LDB1) and to either SBE6 or SBE2 (tSBE6-LDB1 and tSBE2-LDB1) in ESCs (Figure S1D). 3D FISH revealed dramatically increased enhancer-promoter co-localization (≤200nm) upon tShh-LDB1 and tSBE6/tSBE2-LDB1 co-transfection (Figure S1E and S1F). Four-color 3D-FISH, confirmed that a chromatin conformation consistent with a loop, and not simply chromatin compaction, was formed upon Tale-LDB1 expression (Figures S1G and H & Table S2). We conclude that 3D FISH is able to detect a chromatin loop in ESCs, albeit an artificially constructed one.

### Shh-SBE6 chromosome unfolding requires SBE6 and occurs in vivo

We have shown previously that SBE6 is involved in the proper induction of *Shh* expression during NPC differentiation, and that there is a prominent gain of H3K4me1 and H3K27ac at this regulatory element during differentiation to Sox1+ve NPCs (Benabdallah et al., 2016). FISH showed that the increased *Shh*-SBE6 inter-probe distances are abolished in NPCs derived from SBE6^−/-^ ESCs (Figure 2A). This demonstrates that chromatin unfolding across the 100kb 5’ of *Shh* during NPC differentiation is dependent on SBE6.

**Figure 2.**
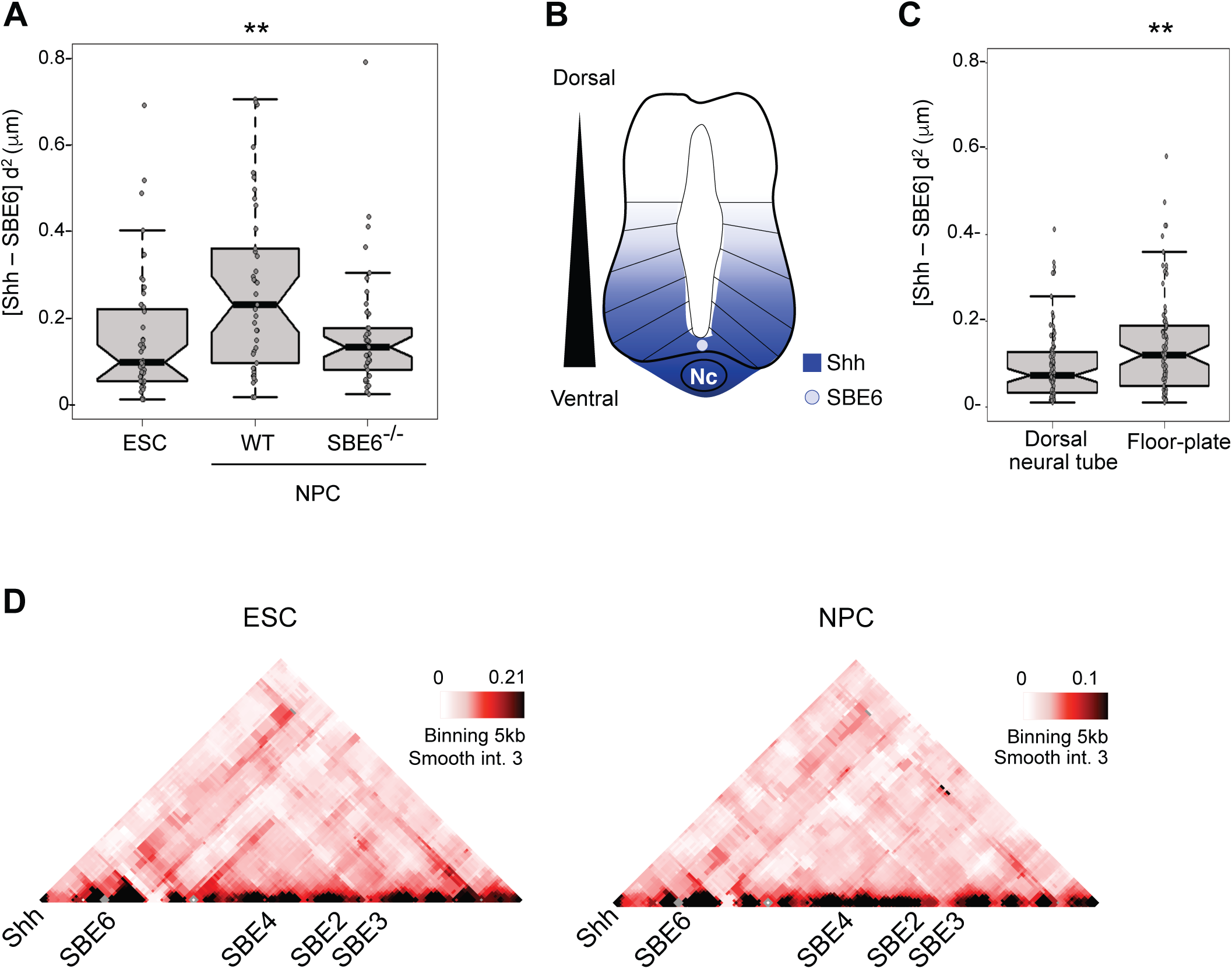
Chromatin unfolding is SBE6 dependent and occurs in vivo. A) Boxplots showing the distribution of squared interprobe distances (μm) between *Shh* and SBE6 in ESCs and in wild-type NPCs as well as in NPCs derived from SBE6.1^−/-^ cells. ** *p*- <0.01. B) Schematic of a transverse section through the neural tube with a gradient of Shh emanating from ventral *Shh* expressing cells in the notochord (Nc) and the floor-plate, where SBE6 activity is also detected (Benabdallah et al., 2016). C) Shh-SBE6 inter-probe distances as in (A) but for FISH data from E10.5 dorsal neural tube and ventral floor-plate cells. ** *p* <0.01 for this biological replicate. For two other biological replicates p = 0.002 and p<0.001. D) 5C heat-maps of the *Shh* regulatory region (mm9, chr5:28750000-29450000) from ESCs and NPCs with 5kb binning and smoothing with the median of 3 surrounding interactions frequencies.

Mouse transgenic assays indicate that SBE6 is active in brain development and in the neural tube (Benabdallah et al., 2016). *Shh* is expressed specifically in ventral regions of the neural tube – in the floorplate and notochord (Jeong et al., 2006) (Figure 2B) and SBE6 drives floorplate expression in a reporter assay. To assess if chromatin unfolding occurs *in vivo*, we used FISH to examine Shh-SBE6 inter-probe distances in sections through the neural tube of an E10.5 embryo. *Shh*-SBE6 distances were greater in nuclei from the floorplate region than in dorsal neural tube cells (Figure 2C), suggesting that our *ex vivo* analysis reflects a long-range chromatin reorganisation 5’ of *Shh* that also occurs *in vivo* during neurogenesis.

To determine if other, as yet unidentified, cis-regulatory elements interact with the *Shh* promoter during the differentiation of ESCs to NPCs, we used Chromosome Conformation Capture Carbon Copy (5C) to assay cross-linked ligation frequencies across the *Shh* regulatory domain. Consistent with published Hi-C data from ESCs (Smallwood and Ren, 2013), and 5C data from E11.5 embryos (Williamson et al., 2016), 5C revealed that in both ESCs and NPCs all of the SBEs are contained in a topologically associated domain (TAD) that extends from downstream of *Shh* to downstream of *Nom1* (Figure S2A). Comparison of 5C data from ESCs and NPCs revealed no evidence for a gain of specific interactions in NPCs that might indicate the formation of a ‘loop’ between *Shh* and its neural enhancers (Figures 2D & S2B).

### SBE, and not direct Shh promoter, activation promotes chromatin unfolding

Supercoiling associated with transcription is known to decondense large chromatin domains (Matsumoto and Hirose, 2004; Naughton et al., 2013), therefore the chromatin unfolding we observe during NPC differentiation could occur as a passive consequence of *Shh* transcription. To determine if this is the case, or if chromatin unfolding depends on *Shh* Brain-Enhancer activation, we designed an enhancer bypass experiment, fusing a Tal effector targeted to the *Shh* promoter (tShh) to four repeats of the small viral acidic protein Vp16 (Vp64) that can strongly activate gene expression (Zhang et al., 2011) (Figure 3A), including in ESCs (Therizols et al., 2014). Expression of tShh-Vp64 in ESCs led to the activation of *Shh* expression to levels similar to those seen in differentiated NPCs (Figure 3B), but without perturbing markers of plurpotency or neuronal differentiation (Figure S3A). However, this synthetic activation of *Shh* in ESCs did not lead to the increased *Shh*-SBE6 inter-probe distances observed during NPC differentiation (Figure 3C). Therefore, chromatin unfolding upstream of *Shh* is not simply a consequence of activating *Shh* expression.

**Figure 3.**
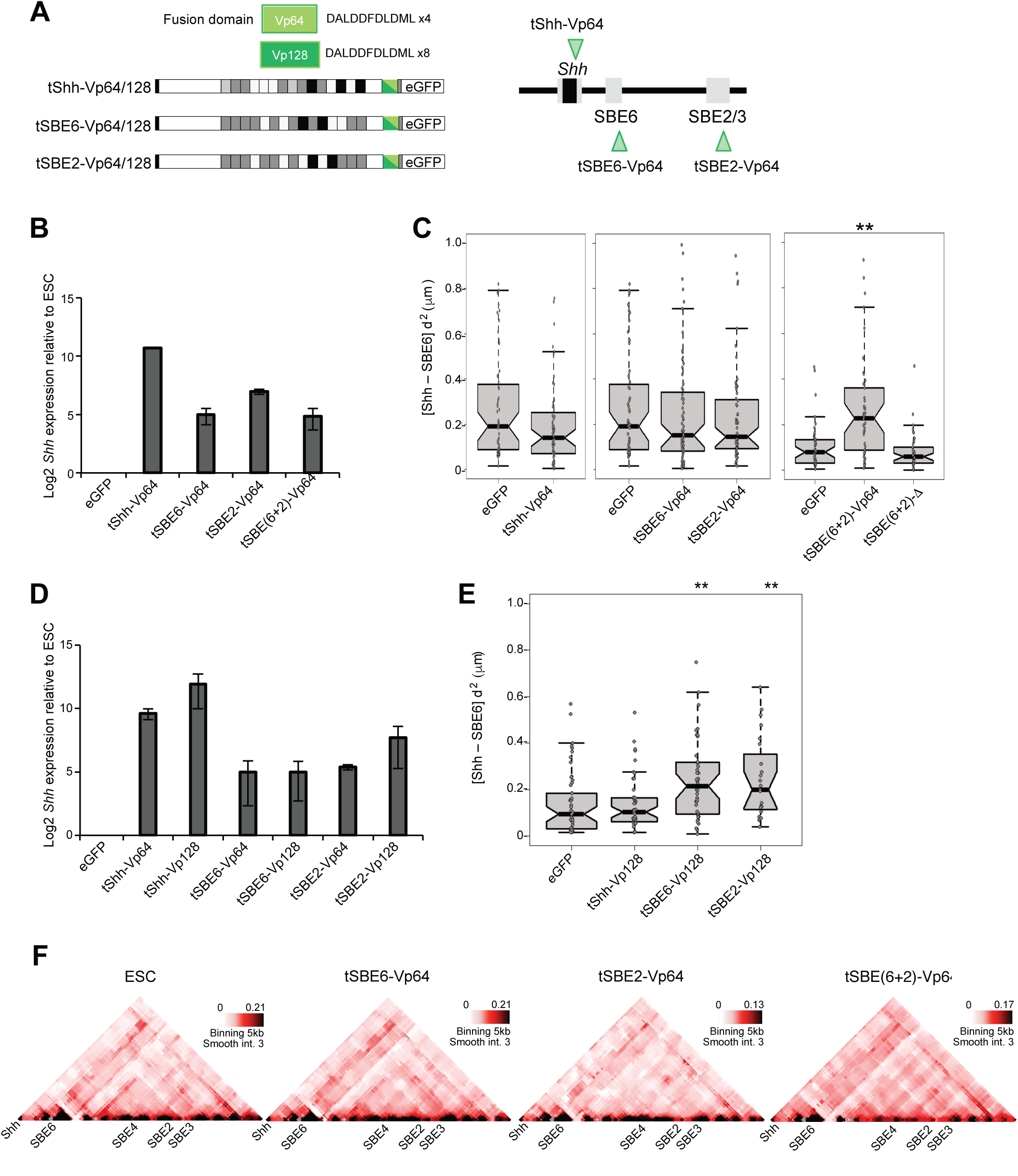
Synthetic activation of Shh and chromatin unfolding using Tale-Vp16. A) Tale-Vp64 and Tales-Vp128 constructs targeting the *Shh* promoter (tShh), SEB6 or SBE2. B) Log2 mRNA levels of *Shh*, relative to *Gapdh*, assayed by qRT-PCR after Tale-Vp64 expression in ESCs. Data show mean (± SEM) of three biological replicates normalized to ESCs expressing a control eGFP. C) Boxplots representing Shh-SBE6 squared interprobe distances (μm) in ESCs expressing; control eGFP, Tale-Vp64 fusions targeting Shh promoter (tShh), SBE6 (tSBE6-Vp64), SBE2 (tSBE2-Vp64), or both SBE6 and SBE2 (tSBE6+2)-Vp64, or a Tal with no fusion protein (tSBE6+2)-Δ. ** p <0.01. D) As in (B) but for both Tale-Vp64 and Tale-Vp128 fusions. E) As in (C) but for Tale-Vp128 fusions. F) 5C heat-maps of the *Shh* regulatory region (chr5:28750000- 29450000) with 5kb binning and smoothing with the median of 3 surrounding interactions frequencies for ESCs and for ESCs expressing Tale-Vp64 fusions targeting SBE6, SBE2 or both SBE6 and SBE2. Statistical analysis for FISH data from this figure and replicate experiments are in Table S3.

Recruiting Vp64 to SBE2 or SBE6 individually, or in combination, through tSBE6-Vp64 and tSBE2-Vp64 also induced *Shh* expression, albeit less marked compared to direct Vp64 recruitment to the *Shh* promoter (Figure 3B). *Shh* activation by Tale-Vp64 recruitment to either SBE2 or SBE6 alone did not lead to increased Shh-SBE6 inter-probe distances (Figure 3C and Table S3). However, simultaneously co-activating SBE6 and SBE2, by co-transfecting both tSBE6-Vp64 and tSBE2-Vp64 (tSBE(6+2)-Vp64), led to significant distance increases between *Shh* and SBE6 (Figure 3C). This was specific to Vp64 activity as recruiting Tales without a fusion domain (tSBE(6+2)-Δ) had no effect (Figure 3C). FISH between Shh and SBE2 confirmed that the chromatin unfolding was confined to the *Shh*-SBE6 region – even when the activator was recruited to SBE2 (Figure S3B and Table S3).

To test whether the differential effect of single vs double Tal-Vp64 targeting was due simply to the local concentration of ‘activator’ that could be recruited, we targeted Vp128 (8 copies of Vp16) to the *Shh* promoter and to SBE6 and SBE2 individually (Figure 3A). All three led to activation of *Shh* expression in ESCs (Figure 3D), but chromatin unfolding was only observed when Vp128 was recruited to the enhancers and not to the *Shh* promoter. In contrast to the recruitment of Vp64, Vp128 recruitment to either SBE6 or SBE2 alone was sufficient to induce this change in chromosome conformation (Figure 3E). Therefore, the amount of activator recruited to the *Shh* regulatory region seems crucial to the induction of visible levels of chromatin unfolding.

5C analysis in ESCs following Tale-Vp64 transfection did not reveal any direct interactions established between the *Shh* promoter and another sequence 5’ of Shh when SBE6 and SBE2 are activated, either individually or together (Figures 3F and S3C).

### Endogenous activators and co-activators also induce chromatin unfolding

Vp16 is a very effective transcriptional activator, but of viral origin. We therefore wished to analyse whether mammalian endogenous activators and co-activators could induce similar long-range chromatin changes across the genomic region 5’ of *Shh*. The Mediator co-activator is known to be recruited to active enhancers and promoters (Allen and Taatjes, 2015; Yin et al., 2014) and can work alongside cohesin to alter 3D chromosome conformation upon enhancer driven gene activation (Kagey et al., 2010; Visel et al., 2009). Chromatin immunoprecipitation (ChIP) showed that Mediator is indeed recruited to the *Shh* promoter during the differentiation of 46c ESCs to NPCs (Figure 4A).

**Figure 4.**
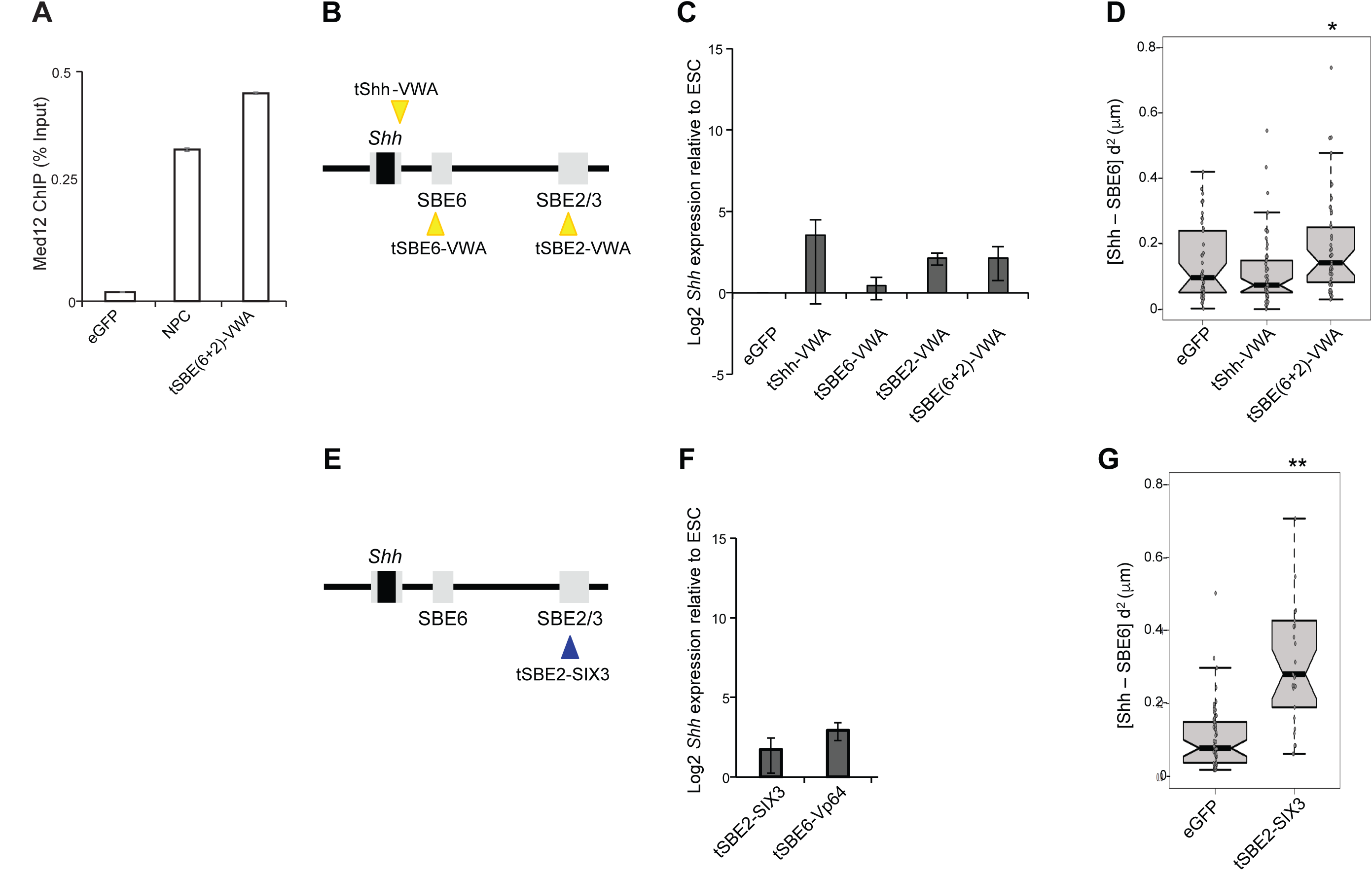
Chromatin unfolding induced by endogenous activators and co-activators. A) Med12 ChIP (% of input, normalised to β-actin promoter) at *Shh* promoter measured by qPCR in NPC or in ESCs expressing eGFP or tSBE(6+2)-VWA. B) Schematic showing the targeting of tale-VWA constructs. C) Log2 mRNA levels of *Shh* relative to *Gapdh* in ESCs expressing Tale-VWA constructs targeting the *Shh* promoter (tShh), SBE6, SBE2, or both SBE6 and SBE2. Data show mean (± SEM) of three biological replicates normalized to ESCs expressing eGFP. D) Boxplots representing Shh-SBE6 squared interprobe distances (μm) in cells expressing eGFP or Tale-VWA targeting the *Shh* promoter (tShh) SBE6, SBE2, or both SBE6 and SBE2. * p = 0.018. E) Schematic showing the targeting of tale-SIX3 construct to SBE2. F) As in (C) but for ESCs expressing tSBE2-SIX3 (13 biological replicates) or tSBE6-Vp64 (11 biological replicates). G) As in (D) but for ESCs expressing tSBE2-SIX3. ** p <0.01. Statistical data relating to FISH data from this figure and replicate experiments are in Table S4.

Mediator interacts with RNA polymerase II and many transcription factors, and is considered to bridge between them. Amongst the transcriptional activators that interact with Mediator is Vp16, which interacts with the Med25 subunit located in the tail domain of the complex (Milbradt et al., 2011; Vojnic et al., 2011). Med25 is recruited into the Mediator complex through its N-terminal Von Willebrand factor A (VWA) domain (Mittler et al., 2003). We therefore fused Tales to the Med25 VWA domain (Figure 4B). Compared to Tale-Vp16 fusions, Med25-VWA recruitment induced only very low levels of *Shh* expression, even for the *Shh* promoter targeted Tale (tShh-VWA) (Figure 4C). Nevertheless, when targeted to both SBE6 and SBE2, Tale-VWA promotes significant chromatin unfolding between SBE6 and *Shh* (Figure 4D and Table S4). Moreover, this also results in the recruitment of the Med 12 subunit of the Mediator kinase module at the *Shh* promoter, compatible with a long-range effect (Figure 4A).

For the most part, the endogenous TFs that bind to SBEs to activate *Shh* in neural tissues are unknown. However, Six homeobox 3 (SIX3) is known to bind to SBE2, and mutation of its binding site, or of SIX3 itself, affects *Shh* expression in the developing brain leading to severe holoprosencephaly (HPE) (Geng et al., 2008; Jeong et al., 2008). *Six3* is also significantly upregulated during the *ex vivo* differentiation of ESCs to NPCs (Benabdallah et al., 2016). This prompted us to investigate whether SIX3 binding was sufficient to recapitulate chromatin unfolding in the region 5’ of *Shh*. Tale-directed recruitment of SIX3 to SBE2 in ESCs (Figure 4E) induced only very low level and variable *Shh* expression (Figure 4F), but led to dramatic chromatin unfolding (Figure 4G, Table S4). Therefore recruitment of either an endogenous activator (SIX3) or co-activator (Mediator) to the gene desert 5’ of *Shh* is capable of inducing an unfolding of long-range chromatin structure and this is not simply dependent on the induction of very high levels of *Shh* expression.

### Blocking of chromatin unfolding suggests a spreading-like mechanism

Chromatin unfolding (increased nuclear distances between SBE6/2 and *Shh*), and the absence of detectable enhancer-promoter looping/spatial juxtaposition, is suggestive of a more linear spreading/linking or tracking-like mechanism operating in the region 5’ of *Shh* (Bulger and Groudine, 1999; Engel et al., 2008; Vernimmen and Bickmore, 2015). To test this model, we attempted to insert obstacles between SBE6 and *Shh* which might block such a mechanism. To do this we chose a site 65kb upstream from the *Shh* TSS (chr5: 28859721; mm9) that lacked evidence of enhancer activity (H3K4me1/H3K27ac marks) during ESC-NPC differentiation (Benabdallh et al., 2016), and that does not show evolutionary conservation. We designed a Tale construct specific to this site (called NE – for Non-Enhancer) and fused it to CTCF (tNE-CTCF) (Figure 5A). CTCF has been proposed to block an enhancer-promoter tracking mechanism at the H19/Igf2 locus (Engel et al., 2008) and to have general enhancer blocking functions (Burgess-Beusse et al., 2002). The introduction of tNE-CTCF, in conjunction with Tale-Vp64 co-activation of SBE6 and SBE2 reduced *Shh* activation (Figure 5B). Single cell qRT-PCR confirmed that the majority of the cells transfected with tNE-CTCF+tSBE(6+2)-Vp64 had very low levels of *Shh* expression (Figure 5C). The binding of tNE-CTCF prevented *Shh*-SBE6 chromatin unfolding induced by the Tale-Vp64 co-activation of SBE6 and SBE2 (Figure 5D and Table S5) consistent with the intervening CTCF molecule interrupting a spreading mechanism initiated at SBE6/2. However, we found that the binding at NE of a Tale without any fused protein (tNE-Δ) had the same negative impact on *Shh* expression (Figure 5B) and *Shh*-SBE6 chromatin unfolding (Figure 5E) as tNE-CTCF. Chromatin unfolding induced by Tale-mediated recruitment of Med25-VWA or SIX3 to SBE6/2 could be similarly blocked by co-transfection with tNE-CTCF or tNE-Δ (Figures 5F and 5G). These data suggest that induced *Shh*-SBE6 chromatin unfolding occurred through a mechanism that is impeded by a protein bound strongly at an intervening site, and that this negatively impacts on *Shh* activation from a distance.

**Figure 5.**
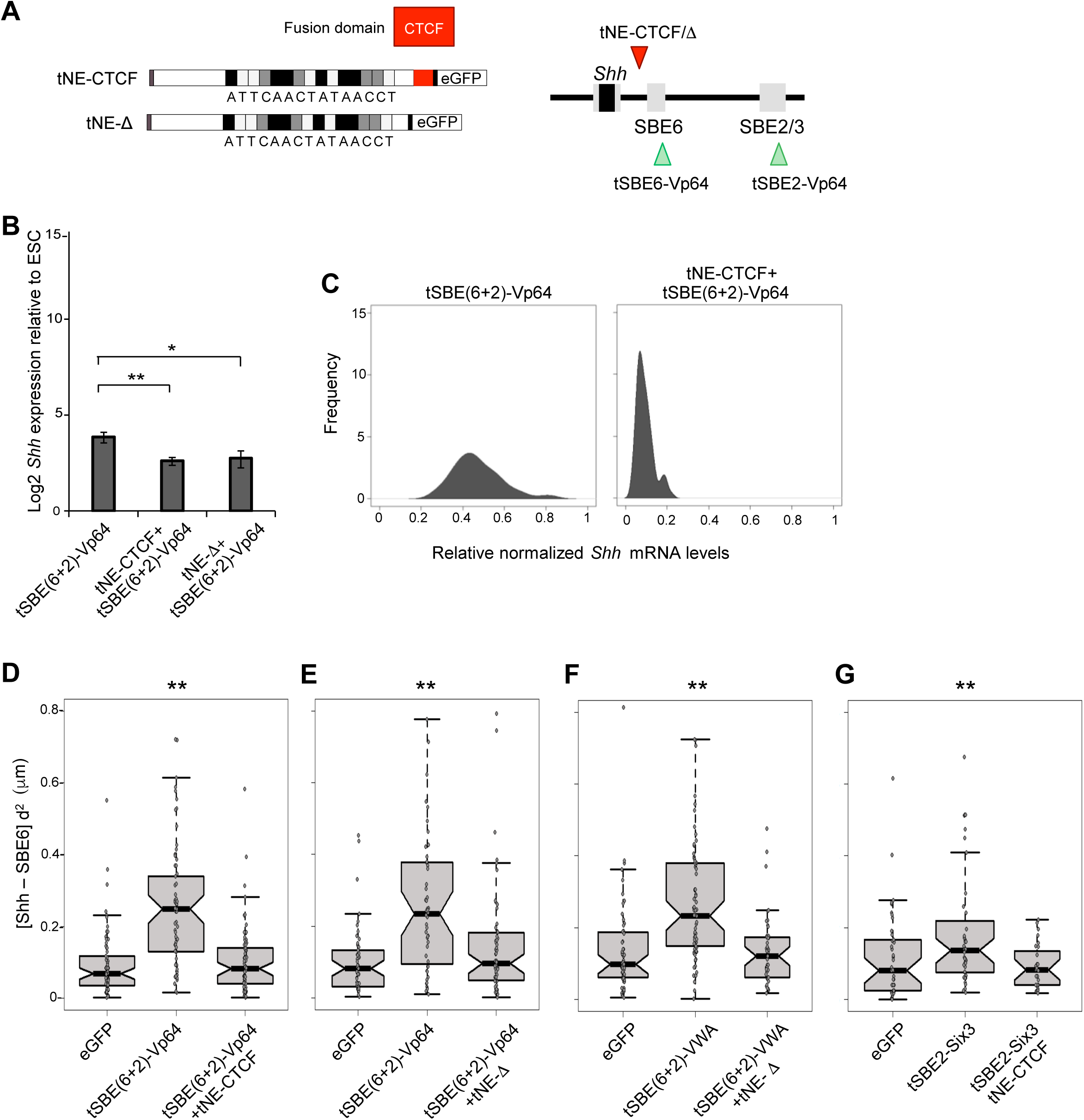
Chromatin unfolding is blocked by an intervening chromatin-bound proteins. A) Schematic showing Tales targeting the NE site with either no fusion protein (tNE-Δ) or fused to CTCF (tNE-CTCF). B) Log2 mean (± SEM) *Shh* mRNA levels relative to *Gapdh* in ESCs expressing Tale-VP64 fusions targeting SBE6 and SBE2 and in cells that also express either tNE-CTCF (3 biological replicates) or tNE-Δ (4 biological replicates). Data are normalized to those from ESCs expressing control eGFP. Asterisks represent p-values for one-tailed Student t-test between conditions. * p <0.05 and ** p <0.01. C) Kernel density plots showing *Shh* expression in single ESCs expressing Tale-VP64 fusions targeting SBE6 + SBE2, and in these cells when tNE-CTCF is also expressed. Expression is normalised to that in ESCs. D and E) Boxplots showing Shh-SBE6 squared interprobe distances (μm) in ESCs expressing eGFP, Tale-VP64 fusions targeting SBE6 + SBE2, and these cells when either tNE-CTCF (D) or tNE-Δ (E) is also expressed. ** p <0.01. F) As in E but using a Tale-VWA fusion targeting SBE6 and SBE2. G) As in (D) but using tSBE2-SIX3 to activate *Shh*. Statistical data for FISH data from this figure arc in Table S5.

### Chromatin unfolding in the Shh regulatory domain involves poly(ADP-ribosyl)ation

Our data suggest that chromatin unfolding spreads within the region 5’ of *Shh*. Histone acetylation can lead to cytological levels of chromatin decompaction (Toth et al., 2004; Lleres et al., 2009) and there are examples of histone acetylation spreading between an enhancer and target gene, and that is blocked by CTCF (Zhao and Dean, 2004). *Shh* activation using Tale-Vp64 constructs targeted to SBE6 or SBE2 did indeed induce histone acetylation (H3K27ac) but this was limited precisely to the Tale binding site – with no indication of spreading (Figure S4).

Another post-translational chromatin modification that can induce large-scale chromatin unfolding *in vitro* and *in vivo* is poly(ADP-ribosyl)ation (PARylation) catalysed by poly(ADP-ribose) polymerases, including PARP1 (Huletsky et al., 1989). Moreover, PARP1 and high levels of poly(ADP-ribose) have been linked with chromatin decompaction and gene activation at ecdysone and heat-shock induced puffs on *Drosophila* polytene chromosomes (Tulin and Spradling, 2003; Sawatsubashi et al., 2004). We therefore investigated whether PARP1 recruitment could induce chromatin unfolding at the *Shh* region, by targeting PARP1 to *Shh*, SBE6 or SBE2 using Tales (Figure 6A). PARP1 recruitment had a minimal affect on *Shh* expression (Figure 6B), but led to chromatin unfolding between *Shh* and SBE6 when targeted to SBE6 or SBE2 (Figure 6C and Table S6). Recruitment of PARP1 to the *Shh* promoter had no detectable affect on chromatin unfolding in the region. Moreover, PARP1 mediated chromatin unfolding, that had been initiated from either SBE6 or SBE2, could be blocked by co-transfection with tNE-CTCF (Figure 6C).

**Figure 6.**
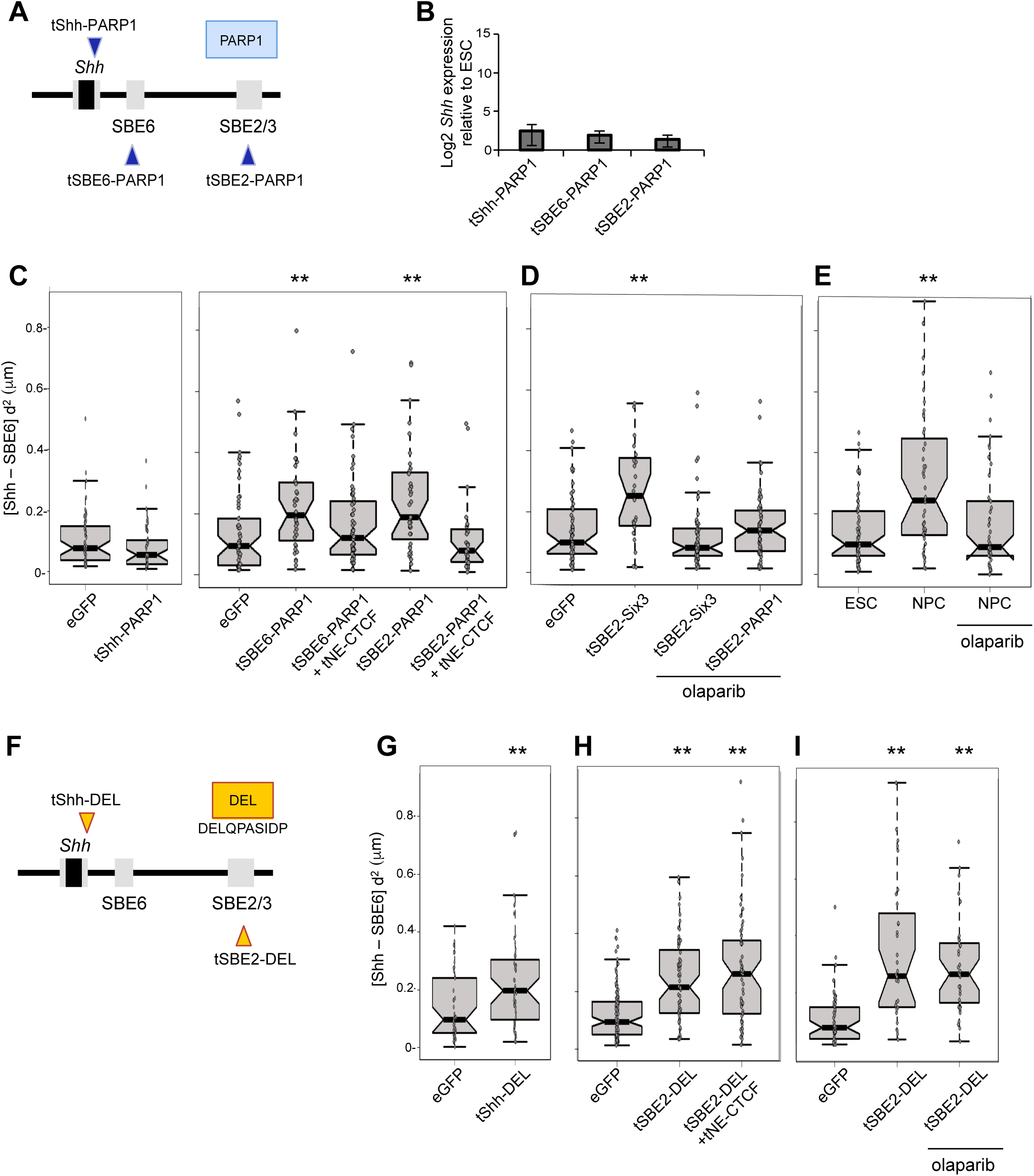
PARP1 catalytic activity induces chromatin unfolding. A) Schematic of Tales that target PARP1 to the promoter (tShh), SBE6 or SBE2. B) Log2 mRNA levels of *Shh* relative to *Gapdh* in ESCs expressing Tale-PARPl constructs targeting the *Shh* promoter (tShh), SBE6, or SBE2. Data show mean (± SEM) of 5 biological replicates normalized to ESCs expressing gGFP. C) Boxplots showing Shh-SBE6 squared interprobe distances (μm) in ESCs expressing; (left) eGFP or tShh-PARP1, (right) eGFP, tSBE6-PARP1, tSBE6-PARP1 + tNE-CTCF, tSBE2-PARP1, tSBE2-PARP1 + tNE-CTCF. ** *p* <0.01. D) As in C but for ESCs expressing eGFP, tSBE2-Six3, and then in tSBE2-Six3 or tSBE2-PARP1 expressing cells treated with olaparib. E) Boxplots showing Shh-SBE6 squared interprobe distances (μm) in ESCs or NPCs, and in NPCs treated with olaparib. F) Schematic of Tales that target the DEL peptide to the *Shh* promoter or to SBE2. G-I) Boxplots showing Shh-SBE6 squared interprobe distances (μm) in ESCs expressing (G) eGFP or tShh-DEL, (H) tSBE2-DEL or tSBE2-DEL + tNE-CTCF, (I) tSBE2-DEL with and without olaparib treatment. Asterisks on FISH data represent Mann-Whitney U test significance between Tale-BP and eGFP transfected or ESC populations, ** for p-values <0.01. Statistical data relating to FISH data are shown in Table S6.

To assess if the chromatin unfolding seen by targeted recruitment of other activators – such as SIX3 (Fig 5C) – also involves PARP1 catalytic activity, we used the PARP inhibitor olaparib (Shen et al., 2015). Olaparib treatment prevented chromatin unfolding mediated by either SIX3 or PARP1 recruitment to SBE2 (tSBE2-SIX3 or tSBE2-PARP1) (Figure 6D). Olaparib also prevented the *Shh*-SBE6 distance increases seen upon the differentiation of ESCs to NPCs (Figure 6E and Table S6).

We have previously shown that large-scale chromatin unfolding can be induced by Tale-mediated recruitment of a small acidic peptide DELQPASIDP (DEL) which has been shown to decompact chromatin without leading to gene activation (Carpenter et al., 2005; Therizols et al., 2014). Targeting DEL to the *Shh* region (Figure 6F) also led to visible chromatin decompaction (Figure 6G and H), but unlike our experiments tethering Vp16, Mediator, or PARP1, this was also seen when the DEL peptide was recruited directly to the *Shh* promoter (Figure 6G). This result is similar to our previous studies in which the DEL peptide was recruited to the promoters of silent genes in ESCs (Therizols et al., 2014). Also unlike the chromatin decompaction induced by PARP1, transcriptional activators and co-activators, chromatin unfolding induced by the DEL peptide recruited to SBE2 was not blocked by tethering of an intervening CTCF (Figure 6H and Table S6), and it was not inhibited by olaparib (Figure 6I). These data suggest that chromatin decompaction induced in the regulatory region 5’ of *Shh*, either during NPC differentiation or during synthetic activation, is quite distinct in its mechanism and its mode of propagation from that induced by the DEL peptide.

To better understand the nature of the unfolded chromatin upstream of *Shh*, we performed 2D-FISH with a fosmid probe for *Shh*, and a BAC probe that spans the 171kb region from 50kb 5’ of *Shh* up until SBE4 (Figure 7A) – the limit of the chromatin unfolding detected during NPC differentiation (Figure 1B). BAC probe hybridisation signals were classified as either compact point, or extended/‘puffed’ signals. Relative to control ESCs, there was a significant increase in the proportion of puffed signals observed when either Vp16 or PARP1 was recruited to the *Shh* regulatory domain, and this was inhibited when PARP1 recruitment was conducted in the presence of olaparib (Figure 7A and Table S7). These data suggest that there is a decompaction of the entire chromatin region extending 200kb 5’ of *Shh* when either an activator (Vp16) or PARP1 is recruited to this region, that requires the catalytic activity of PARP1 and that resembles the puffs seen on Drosophila polytene chromosomes.

**Figure 7.**
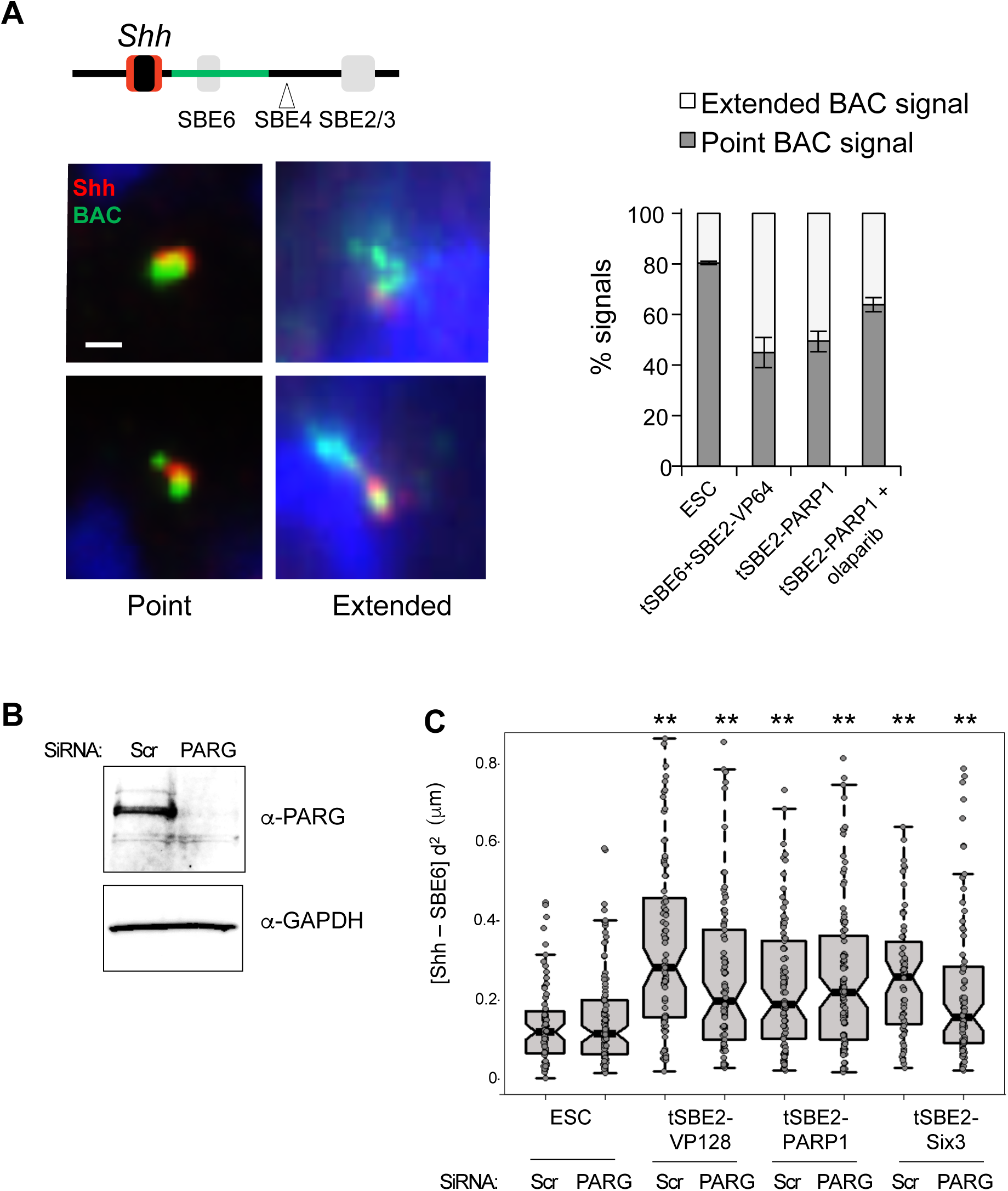
Chromosome puffing induced by activators and PARP1. A) FISH with probes for *Shh* (red) and a BAC probe (RP24-323C22) spanning a 171kb region 5’ of *Shh* (green). Examples of FISH signals with BAC signal classified as either “single point” (left) or extended (right) are shown. The bar chart shows the proportion of ‘single’ vs ‘extended’ BAC signals for ESC, tSBE(6+2)-Vp64, tSBE2-PARP1 and tSBE2-PARP1 treated with olaparib. Data are the average (± SEM) of three technical replicates. Statistical significances from Fishers exact test and total signal counts are displayed in Table S7. B) Immunoblot with antibodies detecting PARG (top) or GAPDH control (bottom) in whole cell extracts from ESCs transfected with scrambled control siRNAs or SiRNAs targeting Parg. D) Boxplots representing Shh-SBE6 squared interprobe distances in ESCs, and in ESCs transfected with tSBE2-Vp128, tSBE2-SIX3, and tSBE2-PARP1 in cells transfected with scrambled (scr) or Parg siRNAs. Asterisks represent Mann-Whitney U test significance between Tales and scramble siRNA transfected ESC, * for p-values < 0.05, ** for p-values <0.01. Statistical data relating to FISH data are shown in Table S8.

Two scenarios can be envisaged for how PARP1 activity and PARylation contribute to large-scale chromatin remodelling. PARylation, e.g. of a chromatin substrate, could lead directly to chromatin decompaction. Alternatively, hydrolysis of the ADP-ribose chains by poly(ADP-ribose) glycohydrolase (PARG) could be being used to generate ATP in the nucleus to support the activity of ATP-dependent chromatin remodelling enzymes (Wright et al., 2016). To distinguish between these two mechanisms, we depleted PARG using siRNA mediated knocked-down (Figure 7B). Relative to ESCs, chromatin decompaction in the *Shh*-SBE6 region, induced by recruitment of Vp16, PARP1 or SIX3, was still detected after PARG knockdown (Figure 7C and Table S8). We conclude that most of the chromatin decompaction that we have detected as a result of recruitment of activators or PARP1 to the regulatory region 5’ of Shh relies on PARP1 but not on PARG activity and so is like due to the process of PARylation itself.

## Discussion

A popular model of enhancer-promoter communication involves chromatin looping to juxtapose the two elements in 3D nuclear space (Vernimmen and Bickmore, 2015), and indeed we have provided visual evidence that supports this model in the context of long-range gene activation of *Shh* by the ZRS enhancer during limb development (Williamson et al., 2016). Recent results using live-cell imaging in *Drosophila melanogaster* to study nascent transcription driven by enhancer elements have challenged the idea of stable enhancer-promoter loops as the basis for enhancer-promoter communication (Fukaya et al., 2016). However, only a handful of enhancer-promoter interactions have been studied in detail and validated, and therefore the generality of a chromatin looping mechanism remains unclear. Here we have analysed the spatial relationship between *Shh* and its neural enhancers SBE6, SBE4 and SBE2/SBE3 using FISH and 5C during neural differentiation and, surprisingly, we identified a visible decompaction of the chromatin in the 350kb region between *Shh* and SBE4, with no evidence for chromatin loops being formed between *Shh* and enhancers in this region. Chromatin unfolding is dependent on a regulatory element (SBE6), that we have previously shown becomes activated during NPC differentiation, and that *in vivo* has activity in the floor plate region of the neural tube – an important site of *Shh* expression (Benabdallah et al., 2016). Consistent with these data, chromatin decompaction between *Shh* and SBE6 is detected in the floor plate, but not in the dorsal neural tube in mouse embryos.

To dissect the mechanism leading to this chromatin decompaction, we used synthetic activators (based upon Tale-mediated recruitment of Vp16) to either induce *Shh* expression directly through activator recruitment to the gene promoter, or to induce expression from a distance through recruitment to distal sites up to 400kb away in the *Shh* regulatory domain. Activation from a distance recapitulated the chromatin decompaction seen during NPC differentiation and this was not seen if activators, or co-activators were recruited directly to the *Shh* promoter, allowing us to exclude that large-scale chromatin decompaction is simply a consequence of *Shh* expression. Induction of chromatin decompaction was recapitulated using distal recruitment of an endogenous activator (Six3) or co-activator (Mediator).

Chromatin decompaction seems most compatible with a tracking or linking mechanism of enhancer action (Bulger and Groudine, 2011). Indeed, we could block chromatin decompaction and abrogate *Shh* activation from a distance if a Tal effector protein was bound between *Shh* and the distal site of activator/coactivator recruitment. This was not generic to all forms of visible chromatin decompaction - for example that induced by recruitment of the DEL peptide (Carpenter et al., 2005; Therizols et al., 2014). Together, our results reveal a strong link between a distinct form of chromatin unfolding and robust *Shh* activation from a distance.

### PARP-dependent chromatin puffing and enhancer activation

Both gene activation from a distance, and chromatin unfolding at *Shh*, could be induced by PARP1 recruitment and blocked by a PARP inhibitor (olaparib). PARylation has been shown to induce massive decompaction of nucleosome arrays *in vitro* (Poirier et al., 1982; Huletsky et al., 1989) and *in vivo* (Tulin and Spradling, 2003; Petesch and Lis, 2008). Substrates for PARylation include the core histones, histone H1 and other proteins that can remodel and relax chromatin structure (Gottschalk et al., 2009; Petesch and Lis, 2008; Timinszky et al., 2009; Ji and Tulin, 2010; Sellou et al., 2016).

Though PARP1 is usually studied in the context of DNA damage sensing and repair, it has also been associated with the regulation of gene expression (Ogino et al., 2007) and with active chromatin and regions with regulatory potential as assayed by DNase I hypersensitivity (Nalabothula et al., 2015). More specifically, PARP and Parp1-dependent PARylation have been implicated in gene regulation from distal enhancer elements that are controlled by nuclear hormone receptors including; ecdysone inducible puffs in Drosophila (Sawatsubashi et al., 2004) and the action of the liganded progesterone receptor in mammalian breast cancer cells (Wright et al., 2012). The function of PARP1 and PARylation at regulatory elements might be directly to open chromatin structure to facilitate access of other factors involved in transcriptional activation. A second possibility is more linked to the role of PARP-1 in DNA repair. There is growing evidence that DNA strand-breaks are important in transcriptional activation. Topoisomerase IIb dependent double-strand break formation, and the subsequent recruitment of Parp-1 activity, is required for signal-induced transcriptional gene activation (Ju et al., 2006). Single-strand nicks generated by topoisomerase I might also be important (Puc et al., 2016). Recruitment of PARP-1 might be through DNA breaks per se, through interaction with transcription factors or co-activator complexes, or through altered DNA structures associated with the activation of transcription (Lonskaya et al., 2005). Finally, the importance of PARylation might be not simply be in decondensing higher-order chromatin structures, but instead the nucleic-acid like PAR chains may seed a phase separation state through liquid de-mixing (Altemeyer et al., 2015) that then organizes an enhancer-centred nuclear compartment that can concentrate or exclude other proteins and orchestrate the multi-step reactions linked to robust transcriptional activation (Hnisz et al., 2017). Further experimentation will be required to investigate these possibilities.

## Experimental Procedures

### Cell Culture, differentiation and transfection

46c mouse embryonic stem cells (mESCs), derived from E14tg2A, contain a GFP insertion into the Sox1 locus (Ying et al., 2003). mESCs were cultured in GMEM supplemented with 10% foetal bovine serum (FBS), 1000 units/ml LIF, nonessential amino acids, sodium pyruvate, 2-β-mercaptoethanol, L-glutamine and penicillin/streptomycin. ESCs were differentiated into NPCs with N2B27 medium (Pollard et al., 2006; Taylor et al., 2013). ESCs were transfected with Tale plasmids using Lipofectamine^®^ 2000 Reagent (Invitrogen cat. N°11668) and FAC-sorted for GFP as previously described (Therizols et al., 2014). Briefly, 1x10^6^ ESCs were transfected in a 6-well plate with 2.5μg of plasmid and 7μl of Lipofectamine. The culture medium was changed 6h after transfection. Transfected cells were sorted based on eGFP expression by FACS 24h after transfection and re-seeded on slides or 6-well-plates. Flow cytometric analysis was performed using the 488nm laser of a BD FACSAriaII SORP (Becton Dickinson) with 525/50 nm bandpass filters. BD FACSDiva software (Becton Dickinson, Version 6.1.2) was used for instrument control and data analysis.

For PARP inhibition, olaparib was added to ESC or NPC media to a concentration of 10μM 24h after transfection or 5h after FACs. For NPCs, cells were treated for 1.5 h and media was changed and cell were fixed for FISH.

For siRNA knockdown, scramble (GE Dharmacon, ON-TARGETplus Non-targeting Pool D-001810-10-05) or PARG (GE Dharmacon, SMARTpool: ON-TARGETplus Parg siRNA L-044091-01-0005) siRNAs were used. 3x10^5^ ESCs were transfected with 100pmol siRNA and 5μl Lipofectamine in a 6-well plate. Media was changed after 16h and at 24h cells were transfected with plasmids as required. Cells were recovered at 72h for expression analysis, immunoblotting and FISH.

### 3D-FISH

1x10^6^ ESCs or NPCs were seeded on slides for 5h. Cells were fixed in 4% paraformaldehyde (pFA) for 10 mins at room temperature (r.t.) and then permeabilized using 0.5% TritonX for 10 mins (Eskeland et al., 2010). Fosmid clones were prepared and labelled with green-dUTP (Abbott Molecular 02N32-050, 00884999002913) or red-dUTP (ChromaTide Alexa Fluor 594-5-dUTP C11400). Approximately 150 ng of labelled fosmid probes were used per slide, together with 15 μg of mouse Cot1 DNA (GIBCO BRL) and 10 μg salmon sperm DNA. Probes were denatured at 70°C for 5 min, reannealed with CotI DNA for 15 min at 37°C and hybridized to the denatured slides overnight. DNA was denatured at 80°C for 20 mins. FISH probes are described in Table S9.

For 2D FISH, cells were swollen in 0.5% trisodium citrate/0.25% KCl followed by fixation in methanol acetic acid (MAA – 3:1 vol/vol). Slides were incubated in 100 μg/ml RNase A in 2 x SCC for 1 hour, washed in 2 x SCC and dehydrated through an alcohol series. Slides were denatured in 70% formamide/2 x SCC for 75 s at 70°C. Between 80-120 ng of biotin- and digoxigenin-labeled probes were used per slide, with 8-12 μg of mouse Cot1 DNA (Invitrogen) and 10 μg salmon sperm DNA. Probes were denatured at 70°C for 5 mins, reannealed with Cot1 DNA for 15 mins at 37°C and hybridized to the denatured slides overnight at 37°C. Slides were washed 4 x 3 minutes in 2X SSC at 45°C, 4 x 3 mins in 0.1X SSC at 60°C and transferred to 4X SCC, 0.1% Tween 20. Slides were counterstained in 0.5 μg/ml DAPI.

### Image Capture and Analysis

Super-resolution images from 3D FISH were acquired using structured illumination microscopy (SIM) following a published protocol (Gustafsson et al., 2008). Samples were prepared on high precision cover-glass (Zeiss, Germany). 3D-SIM images were acquired on a N-SIM (Nikon Instruments, UK) using a 100x Nikon Plan Apo TIRF objective (NA 1.49, oil immersion) and refractive index matched immersion oil (Nikon Instruments). Images were captured using an Andor DU-897X-5254 EMCCD camera using 405, 488, 561 and 640nm laser lines. Step size for *z* stacks was set to 0.12 μm as required by the manufacturer’s software. For each focal plane, 15 images (5 phases, 3 angles) were captured with the NIS-Elements software. SIM image processing, reconstruction and analysis were carried out using the N-SIM module of the NIS-Element Advanced Research software. Images were reconstructed using NiS Elements software (Nikon Instruments) from a *z*-stack comprising of no less than 1μm of optical sections. In all SIM image reconstructions the Wiener and Apodization filter parameters were kept constant.

For analysis of 2D FISH, slides were imaged using a Hamamatsu Orca AG CCD camera (Hamamatsu Photonics (UK) Ltd, Welwyn Garden City, UK) and a Zeiss Axioplan II fluorescence microscope with Plan-neofluar objectives, a 100W Hg source (Carl Zeiss, Welwyn Garden City, UK) and Chroma #83000 triple band pass filter set (Chroma Technology Corp., Rockingham, VT) with the excitation filters installed in a motorised filter wheel (Prior Scientific Instruments, Cambridge, UK).

Image analysis was carried out using the Quantitation module of Volocity (PerkinElmer). Reconstructed SIM data was directly uploaded and analyzed on Volocity. The statistical significance of differences in mean-squared interprobe distances was assessed using the nonparametric Mann-Whitney U test to examine the null hypothesis. Each data set consisted of 20 to 50 nuclei (40 to 100 loci). Biological replicates are shown under their p-values in Supplemental figures and tables.

### RNA extraction and Real Time quantitative Polymerase Chain Reaction (qRT-PCR)

RNA was prepared using RNeasy mini kit (Qiagen) according to the manufacturer’s protocol, including a DNaseI (Qiagen) treatment for 15 mins at r.t. cDNA was synthesized from 2 μg purified RNA with Superscript II reverse transcriptase (Invitrogen) primed with random hexamers (Promega).

Real-time PCR was carried on the Roche LightCycler 480 Real-Time PCR System using a Lightcycler 480 Sybr Green detection kit (Roche). The real-time thermal cycler was programmed as follows: 15 min Hotstart; 44 PCR cycles (95°C for 15 s, 55°C for 30 s, 72°C for 30 s). For transcript analysis, a standard curve for each primer set was obtained using a mix of each of the cDNAs. The relative expression of each sample was measured by the Lightcycler software and normalized to the mean for *Gapdh* from replicates. Finally, the log2 of the ratio relative to eGFP transfected ESCs was calculated when mentioned. Primers for qRT-PCR are listed in Table S10. *Ptn* and *Nrp1* expression primers were taken from (Therizols et al., 2014).

### Single Cell RT-qPCR

RNA reverse transcription and cDNA pre-amplification from single cells were performed as previously described (Dalerba et al., 2011) with some modifications. Each well of a 96-well PCR plate was loaded with 5 μl 2x Reaction Mix, 0.2 μl Superscript III RT/Platinum Taq Mix with RNaseOUT Ribonuclease Inhibitor (Invitrogen Cells Direct One-Step qRT-PCR kit, Life Technologies), 2.5 μl primer mix (containing 200 nM of each gene-specific primer), 1.3 μl H_2_O. Single-cell suspensions from NPC differentiation or transfection were sorted on their GFP reporter into separate wells of the 96-well PCR plate. 32 cells were sorted into one well, to be used for serial dilution for generation of qRT-PCR standard curves. RNA reverse transcription and 22 cycles of cDNA pre-amplification were performed as previously described (Dalerba et al., 2011). The cDNA was diluted 1:5 in H2O and qRT-PCR performed as above using 9 μl of this diluted cDNA.

### TALE Design & Assembly

Tales were designed using TAL Effector Nucleotide Targeter 2.0 software (Doyle et al., 2012) and the assembly was performed following the protocol described in (Therizols et al., 2014). Briefly, TALE DNA binding domains specific to the Shh promoter, SBE6, SBE2, and NE were assembled following the methods described in (Ding et al., 2013). DNA binding domains specific for 16 nucleotide sequences were generated by the modular assembly of 4 pre-assembled multimeric TALE repeat modules (three 4-mer and one 3-mer) into a modified TALEN backbone in which the BamHI-BsrGI fragment containing hFokI2-2A-eGFP was replaced by a gBlocks^®^ (IDT) fragment encoding Vp64-2A-eGFP (Therizols et al., 2014). The BamHI-BglII fragment containing Vp64 of the Tale-Vp64 plasmid was deleted to generate Tale-Δ.

To generate other Tal-fusions, the BamHI-NheI cloning site was further used to fuse the Tale with; the SA domain of LDB1 from gBlocks^®^ (IDT), with CTCF, Six3 or PARP1 from GeneArt^®^ (Life Technologies). To generate Tale-BP, the BamHI-NheI fragment containing Vp64 was replaced by a double strand oligonucleotide encoding the DELQPASIDP peptide (Carpenter et al., 2005; Therizols et al., 2014).

### Chromatin Immunoprecipitation and Microarray processing

For cross-link ChIP, 0.5-1x10^7^ ESC were first cross-linked with EGS (Pearce, Thermo Scientific, product no. 21565) in PBS at a final concentration of 2 mM for 60 min at r.t. Formaldehyde was then added at a final concentration of 1% methanol-free formaldehyde (Thermo Scientific Pierce PN28906) and incubated at r.t. for 10 min followed by 5 min incubation with 125mM glycine and then washed in PBS. All buffers were supplemented with the following additives just prior to use: 0.2 mM PMSF, 1 mM DTT, 1x Protease inhibitors (Calbiochem, 539134-1SET) and 1x phosphatase inhibitors (Roche, PhosSTOP, 04906837001). Purified DNA was isolated using a QIAquick PCR Purification Kit (Qiagen). Med12 ChIP was performed as previously described (Vernimmen et al., 2007) using Med12 antibody (Bethyl laboratories, A300-774A) and quantitative (q)PCR was performed on a LightCycler480 (Roche) using the same guideline as for qRT-PCR. ChIP qPCR primers are displayed in Table S10.

For examination of histone acetylation by native ChIP, nuclei were prepared and resuspended in NB-R as previously described (Gilbert et al., 2003). Nuclei corresponding to 0.5-1x10***^7^*** ESCs were digested with 50-80 Boehringer units of MNase (Sigma) for 10 min at r.t. in the presence of 20 μg RNase A to obtain a chromatin ladder enriched in tri-, tetra-, and some pentanucleosomes. The reaction was stopped by adding equal volume of Stop Buffer (215 mM NaCl, 10 mM TrisHCl pH 8, 20 mM EDTA, 5.5 % Sucrose, 2 % Triton X-100, 0.2 mM PMSF 1 mM DTT and complete protease inhibitor cocktail) and incubated on ice overnight. Between 50-150 μg released chromatin were precleared with Protein G Sepharose (GE Healthcare) for 2 hr and mixed with 10 μg prebound H3K27ac antibody (Millipore 07-360) in the presence of 100 μg BSA and incubated for 3 hr at 4 °C. Beads were then washed 3x with Wash Buffer (150 mM NaCl, 10 mM TrisHCl pH 8, 2 mM EDTA, 1% NP40, 1 % Sodium deoxycholate, 0.2mM PMSF, 1mM DTT and protease inhibitor cocktail) and once in TE. Bound complexes were eluted with 0.1 M NaHCO3, 1 % SDS at r.t. Immunoprecipitated and input DNA were purified with Proteinase K (Genaxxon) and Qiagen PCR purification kit.

For Nimblegen Arrays (H3K27ac), 10ng of input (MNase digested) or ChIP DNA were amplified using the WGA2 whole genome amplification kit (Sigma). Amplified material was labelled with Cy3 or Cy5 by random priming according to the NimbleGen ChIP-chip protocol (Roche). Samples were hybridized for 20 h and washed according to manufacturer’s protocol. A custom 3x720K mouse tiling array (NimbleGen, Roche) containing 179,493 unique probes from different genomic regions, with each probe represented by 4 replicates was used. Arrays were scanned on a NimbleGen MS 200 Microarray scanner (Roche) using 100% laser power and 2 μm resolution. Raw signal intensities were quantified from TIFF images using MS 200 Data Collection software.

ChIP microarray data were analysed in R using the bioconductor packages Beadarray and Limma according to the Epigenesys NimbleGen ChIP-on-chip protocol 43. Scale normalization was used within replicates, to control inter-array variability. Enrichment scores are defined as log2 ChIP/Input signal.

### 5C primer, 5C library design and preparation

3C library preparation was performed as previously described (Williamson et al., 2014). 1 x 10^7^ ESCs or NPCs were fixed with 1% formaldehyde for 10 min at r.t. Cross-linking was stopped with 125 mM glycine for 5 min at r.t. followed by 15 min on ice. Cells were centrifuged at 400 *g* for 10 min at 4°C, supernatants were removed, and cell pellets were flash-frozen on dry ice. Cell pellets were then treated as previously described (Dostie and Dekker, 2007; Ferraiuolo et al., 2010; Williamson et al. 2016).

Briefly, 1-2 x 10^7^ fixed cells were incubated for 15 min on ice in 200 μl of lysis buffer (10 mM Tris at pH 8.0, 10 mM NaCl, 0.2% NP40, supplemented with fresh protease inhibitor cocktail). Cells were then disrupted on ice with a dounce homogenizer (pestle B; 2 × 20 strokes); cell suspensions were transferred to Eppendorf tubes and centrifuged at 2000 *g* for 5 min. Supernatants were removed, the cell pellets were washed twice with 100 μl of 1× CutSmart buffer (New England Biolabs), and the cell pellet was resuspended in 100 μl of 1× CutSmart buffer and divided into two Eppendorf tubes. 1× CutSmart buffer (337 μL) was added to each tube, and the mixture was incubated for 10 min at 65°C with 0.1% SDS. Forty-four microliters of 10% Triton X-100 were added before overnight digestion with 400 U of HindIII. The restriction enzyme was then inactivated by adding 86 μl 10% SDS and incubation for 30 min at 65°C. Samples were then individually diluted into 7.62 ml of ligation mix (750 μl 10% Triton X-100, 750 μl 10× ligation buffer, 80 μl 10 mg/ml of BSA, 80 μl 100 mM ATP, 3000 cohesive end units of T4 DNA ligase) and incubated for 2 h at 16°C.

3C libraries were incubated overnight at 65°C with 50 μl of Proteinase K (10 mg/ml) and an additional 50 μl of Proteinase K the following day for 2 h. The DNA was purified by one phenol and one phenol–chloroform extraction and precipitated with 0.1 vol (800 μl) of 3 M NaOAc (pH 5.2) and 2.5 vol of cold EtOH (20 ml). After at least 1 h at −80°C, the DNA was centrifuged at 20,000 *g* for 25 min at 4°C, and the pellets were washed with cold 70% EtOH. DNA was resuspended in 400 μl of TE (pH 8.0) and transferred to Eppendorf tubes for another phenol–chloroform extraction and precipitation with 40 μl of 3 M NaOAc (pH 5.2) and 1.1 ml of cold EtOH. DNA was recovered by centrifugation and washed eight times with cold 70% EtOH. Pellets were then dissolved in 100 μl of TE (pH 8.0) and incubated with1 μl of 10 mg/ml RNase A for 15 min at 37°C.

5C primers covering the USP22 (mm9, chr11: 60,917,307–61,017,307) and Shh (mm9, chr5: 28317087-30005000) regions, library design, and preparation, were performed as described (Williamson et al. 2016). 5C libraries were prepared and amplified with the A-key and P1-key primers as described previously (Fraser et al., 2012). 3C libraries were first titrated by PCR for quality control (single band, absence of primer dimers, etc.) and to verify that contacts were amplified at frequencies similar to that usually obtained from comparable libraries (same DNA amount from the same species and karyotype) (Dostie and Dekker, 2007; Fraser et al., 2010). We used 1–11 μg of 3C library per 5C ligation reaction.

5C primer stocks (20 μM) were diluted individually in water on ice and mixed to a final concentration of 0.002 μM. Mixed diluted primers (1.7 μl) were combined with 1 μl of annealing buffer (10× NEBuffer 4, New England Biolabs) on ice in reaction tubes. Salmon testis DNA (1.5 μg) was added to each tube, followed by the 3C libraries and water to a final volume of 10 μl. Samples were denatured for 5 min at 95°C and annealed for 16 h at 48°C. Ligation with 10 U of Taq DNA ligase was performed for 1 h at 48°C. One-tenth (3 μl) of each ligation was then PCR-amplified individually with primers against the A-key and P1-key primer tails. We used 26 or 28 cycles based on dilution series showing linear PCR amplification within that cycle range. The products from two to four PCR reactions were pooled before purifying the DNA on MinElute columns (Qiagen).

5C libraries were quantified on agarose gels and diluted to 0.0534 ng/μL (for Xpress template kit version 2.0) or 12 pmol (for Ion Proton). One microliter of diluted 5C library was used for sequencing with an Ion Proton sequencer. Samples were sequenced as recommended by the manufacturer (Life Technologies).

Analysis of the 5C sequencing data was performed as described earlier (Berlivet et al., 2013). The sequencing data were processed through a Torrent 5C data transformation pipeline on Galaxy (https://main.g2.bx.psu.edu). Data were normalized by dividing the number of reads of each 5C contact by the total number of reads from the corresponding sequence run. All scales shown correspond to this ratio multiplied by 10^3^. Sequencing technical and biological replicates reads are displayed in Table S11.

### Immunoblotting

3 wells of a 6-well plate containing ESCs transfected with scramble or PARG siRNA were recovered 72h after transfection. Cells were trypsinized, washed and resuspended in 50μl RIPA buffer (10 mM Tris-Cl (pH 8.0) 1 mM EDTA. 1% Triton X-100. 0.1% sodium deoxycholate. 0.1% SDS. 140 mM NaCl. 1 mM PMSF) and left on ice for 30 min. After a 10 min spin at full speed, supernatant was mixed with 1x LDS NuPage loading buffer and 1x reducing agent and boiled at 95°C for 5 minutes. 20μl sample were run in a NuPAGE Novex 3-8% Tris-Acetate Protein Gel (Thermo Fisher Scientific) and transferred using an iBlot^®^ 2 transfer system. Membrane was blocked with 5% Milk in PBS-T for 1h at r.t. For PARG detection, the membrane was incubated for 4 h at r.t. with anti-PARG antibody (PARG M-13, Santa Cruz sc-21480). For the loading control, GAPDH ab9485 was used.

## Authors contributions

NSB conducted most of the experiments and data analysis, prepared the figures and wrote the manuscript. IW assisted with 5C experiments and manuscript editing, RI assisted with ChIP and microarray analysis and manuscript editing, SB assisted with tissue section FISH and 2D FISH, GG assisted the microarray data processing. PT assisted with Tale design and building. WAB conceived of the study, designed the study, coordinated the study and wrote the manuscript. All authors gave final approval for publication. The authors declare that they have no competing interests

## Acknowledgements

We thank the staff of the IGMM FACS and imaging facilities for invaluable technical help.

## Accession numbers

Data from this paper are available at NCBI GEO under the series: GSE89557.

GSE89388: 5C data.

GSE89512: ChIP data

